# Quantum mechanical electronic and geometric parameters for DNA k-mers as features for machine learning

**DOI:** 10.1101/2023.01.25.525597

**Authors:** Kairi Masuda, Adib A. Abdullah, Aleksandr B. Sahakyan

## Abstract

With the development of advanced predictive modelling techniques, we are witnessing a steep increase in model development initiatives in genomics that employ high-end machine learning methodologies. Of particular interest are models that predict certain genomic or biological characteristics based solely on DNA sequence information. These models, however, treat the DNA sequence as a mere collection of four, A, T, G and C, letters, thus dismissing the past physico-chemical advancements in science that can enable the use of more intricate information about nucleic acid sequences. Here, we provide a comprehensive database of quantum mechanical and geometric features for all the permutations of 7-meric DNA in their representative B, A and Z conformations. The database is generated by employing the applicable high-cost and time-consuming quantum mechanical methodologies. This can thus make it seamless to associate a wealth of novel molecular features to any DNA sequence, by scanning it with a matching k-meric window and pulling the pre-computed values from our database for further use in modelling. We demonstrate the usefulness of our deposited features through their exclusive use in developing a model for A to C mutation rate constants.

## Background & Summary

Machine learning techniques are now being actively pursued in all fields, including genomics. The main driver of this phenomenon is the rapid development in computer performance and efficiency as predicted by Moore’s Law^1^. The explosion of digital data necessary to feed the machine learning algorithms has also been a major contributor to the ever-increasing adoption of machine learning. In the field of genomics, big data, especially from high-throughput sequencing technologies, have been utilised to develop many successful machine-learning based models to address various biological problems^2^. Models were built to predict G-quadruplex formation^3^, effective gene expression^4^, splicing events^5,6^, specificities of DNA- and RNA-binding proteins^7^, effects of non-coding variants^8^, epigenomic profiles^9^, transcription factor binding^10^, regulatory code of the accessible genome^11^, DNA methylation^12^, cancer driver genes^13^, and many other biological phenomena.

Most of these works mainly focus on the underlying oligonucleotide letter strings to devise the features for machine learning, requiring a massive amount of data to decipher and exhaust information out of nucleic acid sequences. These sequence-based initiatives, despite being successful, still overlook a decades worth of information and advancement accumulated on the inference of physico-chemical properties of oligonucleotides in their varying sequence context and structure. Many such properties are commonly calculable from molecular modelling, at a coarse and atomistic scales *via* molecular mechanics (MM) and quantum mechanical (QM) electronic techniques. If pre-calculated for different sequence contexts, those properties can be parameterised and seamlessly incorporated in feature generation stage of machine learning, still keeping the model purely sequence-based in terms of the required primary information.

To make the above possible, here we performed a large-scale semi-empirical QM calculations for all possible DNA heptamers in their three, B, A and Z, representative conformations (**Fig. 1**). We began from DNA 3D model development and geometry optimisation, followed by the semi-empirical calculations. A number of geometric values of the optimised models were also measured and included as part of the database. The outcomes of these calculations can be applied as approximate values to the machine learning algorithms, where their relevance and information content will be assessed accordingly during the learning process. As a proof of principle, we briefly demonstrated that our DNA heptamer semi-empirical properties along with their geometric measurements were able to predict A to C spontaneous mutation rates^14^ from DNA sequence, when they were applied as sole machine learning features with no direct sequence encoding.

**Figure 1.**
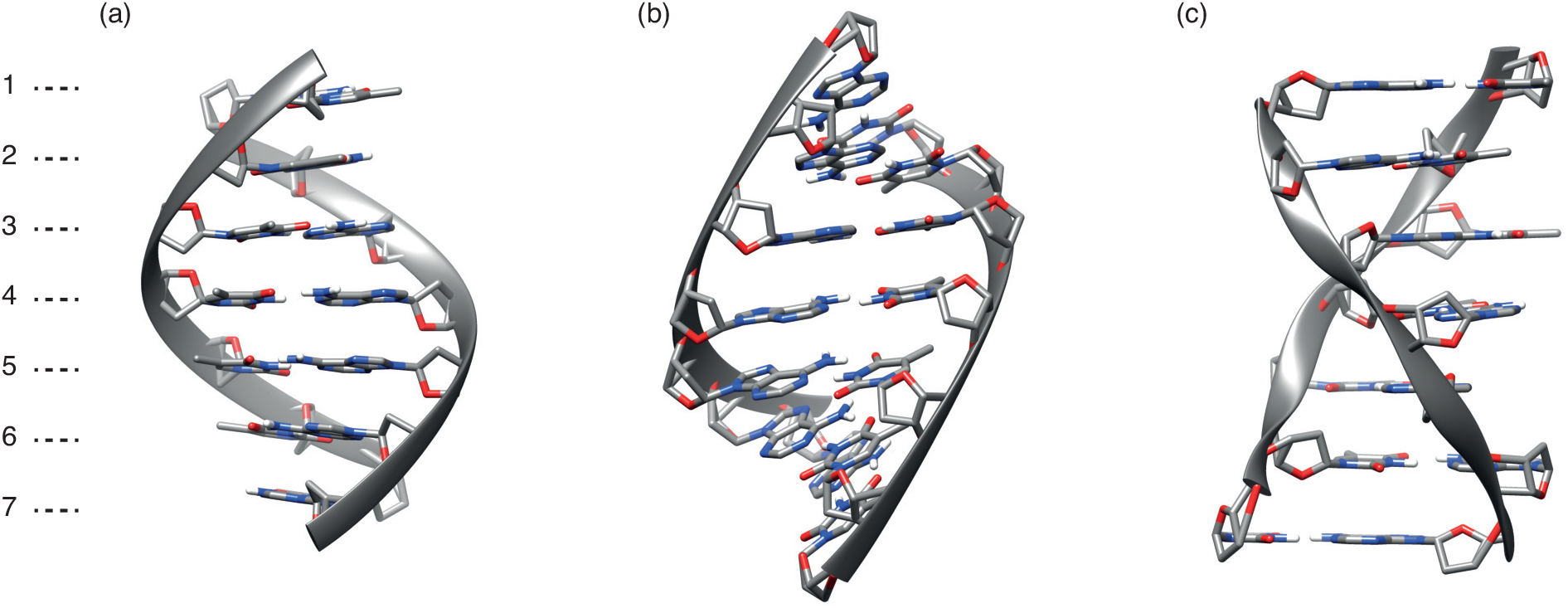
Representative molecular structures of the 7-mer double-stranded DNA models used in this study. The structures are brought for (a) B-DNA, (b) A-DNA, and (c) Z-DNA conformations. For clarity, backbones are represented by ribbons. These structures have seven nucleotide spans at both strands.

We believe our deposited data presented in this work, comprised of 3D geometric and physico-chemical properties of 24,576 non-redundant DNA 7-mer duplex structures (not counting the shorter-span 6-mer analogues) in B, A and Z conformations (**Fig. 1**), will be useful and applicable to enhance the machine learning process in solving DNA-driven biological problems. Such a database is unique for varying oligonucleotides, even though databases of many MM and QM properties exist for other types of molecules, many reported in the same, Scientific Data, journal. Examples of other QM-based datasets published in the same spirit are physico-chemical properties of 31,618 electroactive molecules for development of aqueous redox flow batteries^15^, optimised molecular geometries and thermodynamic data of more than 665,000 biologically and pharmacologically relevant molecules^16^, electronic charge density of crystalline materials from Materials Project database^17^, molecular conformations of 450,000 small- and mid-sized organic molecules^18^, molecular geometries and spectral properties of 61,489 crystal-forming organic molecules^19^, equilibrium conformations for small organic molecules^20^, QM calculations of over 200,000 organic radical species and 40,000 associated closed-shell molecules^21^, all-atom force-field parameters, molecular dynamics trajectories, QM properties, and curated physicochemical descriptors of more than 300 antimicrobial compounds^22^, excited state information of 173,000 organic molecules^23^, conformational energies and geometries of di- and tripeptides ^24^, and QM structures and properties of 134,000 small organic molecules^25^.

## Methods

The schematic diagram on the generation of the database is brought in **Fig. 2**. The procedure is comprised of three stages: the building of the all-atom DNA models (a), geometry optimisation (b), and feature extraction with the corresponding single-point calculations (c). In the following subsections, we describe the details of each stage. The calculations were performed on the available Linux computing clusters hosted at the MRC Weatherall Institute of Molecular Medicine, University of Oxford (256 GB of RAM, dual Intel Xeon E5-2680v3 CPUs with 24 physical cores per node), and on our laboratory workstation (512 GB of RAM, Intel Xeon W-2295 CPUs with 18 physical cores). The database can be accessed through our GitHub page at https://github.com/SahakyanLab/DNAkmerQM. R^26^ was used as a front-end programming language in this work, and the code to generate the database can be retrieved from https://github.com/SahakyanLab/NucleicAcidsQM.

**Figure 2.**
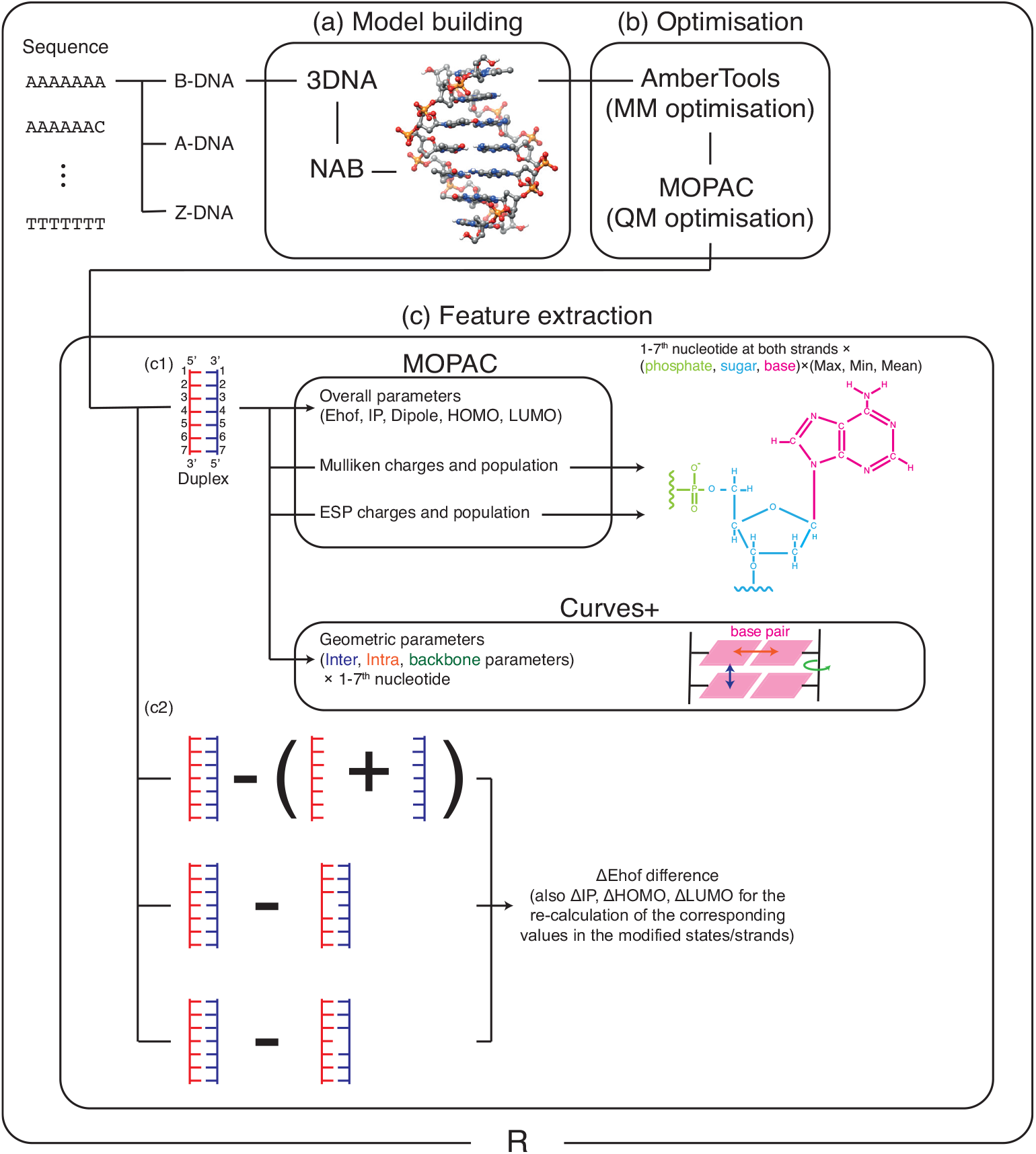
A schematic workflow of the database generation used in this study. In the whole procedure, we used R programming language for processing data. (a) To make molecular DNA models, we used web 3DNA and NAB program, and a typical resultant DNA model is shown as an illustration. (b) Next, we optimised these models in two steps. First *via* MM optimisation by AmberTools21, next *via* QM optimisation by MOPAC2016. (c1) After optimisation, we extracted electronic features by single-point MOPAC2016 calculations and geometric features by using Curves+. For charges and populations, we calculated the maximum, minimum, and mean values of phosphate, sugar, and base moieties. (c2) We further conducted single-point calculations for each strand separately, and for the states with deleted central base. Next, the differences in energy and overall features between the duplex and these states were calculated. Note, that differences in IP, HOMO, and LUMO (ΔIP, ΔHOMO, and ΔLUMO) from varying molecular states do not have clear physical meaning and should be avoided from direct usage in machine learning. However, we include them in our database to enable the retrieval of the corresponding values for single-stranded and base-deleted states from the IP, HOMO, and LUMO of duplex states, i.e. by taking IP-ΔIP, HOMO-ΔHOMO and LUMO-ΔLUMO.

### All-atom model building for 7-mer DNA

As the maximum context-span and the baseline in this work, heptameric range of DNA was considered due to its known major influence on nucleotide and derivative properties^14^. However, our database also includes an analogue for the lesser, hexameric context, mainly generated for the use cases when even-numbered range is needed for modelling. The DNA structures for all the k-mer sequence permutations in their B, A and Z conformations (**Fig. 1**) were generated by using the Nucleic Acid Builder (NAB) suit of programs^27^ (**Fig. 2a**). NAB provides a function that replaces base pairs of a given template structure with any desired base pair, without altering the geometries of the backbone and sugar moieties. As templates for B, A and Z conformations of double-stranded DNA, representative X-ray crystallographic structures^28^ were used as adapted from the PDB files provided in WEB-3DNA^29^ (**Fig. 2a**). The end moieties for the 1^*st*^ and 7^*th*^ positions in the DNA models were capped by hydrogen atoms (at the O5’ and at O3’ positions of deoxyribose for the 5’ and 3’ ends respectively). The total number of all the permutations for 7-mer sequences of four bases is 4^7^ =16,384. However, considering the strand symmetry of double-stranded DNA, we can reduce this number, since, for instance, the sequence 5’-AAAAATT-3’ has 5’-AATTTTT-3’ as a complementary strand, hence the double-stranded DNA model for 5’-AAAAATT-3’ is the same for 5’-AATTTTT-3’ as well. We therefore generated 8,192 DNA models for each B, A and Z conformations, resulting into a total of 24,576 DNA models for heptamers. On average, one DNA model in our generated set contains 443 atoms (281 heavy, and 162 hydrogen atoms).

### Molecular mechanics optimisation of DNA

The above DNA structures were then geometry optimised *via* molecular mechanics (MM) force field, by using AMBER-TOOLS21^30^ (**Fig. 2b**). The OL15 force field, specifically tuned and well tested for DNA^31^, was used. To account for the shielding of the negatively charged phosphate backbones, Born implicit solvation^32^ was used with the water environment dielectric constant defaulted to 78.5 in AMBERTOOLS21. We needed to relax the geometries in order to remove any tension and unrealistic arrangements upon NAB-driven base replacements. However, we still wanted to preserve the conformations in their desired original, B, A or Z, state. We therefore had to pick an appropriate number for the allowed optimisation steps. **Fig. S1a1** shows a relationship between root mean squared deviation (RMSD) of 5’-AAAAAAA-3’ B-DNA, from its original NAB-generated structure, and the number of optimisation steps. RMSD calculation was done using the BIO3D library^33^ in R, based only on non-hydrogen atoms. We found that RMSD converged within 5000 steps. Furthermore, no major conformational change or strand separation was observed in the structures before and after the convergence (**Fig. S1a2**). **Fig. S1a3** shows the RMSDs for all the heptamers of B-DNA. Note that the sequences are numbered lexicographically, that is, 5’-AAAAAAA-3’=1, 5’-AAAAAAC-3’=2, 5’-AAAAAAG-3’=3, and so on. The results show that the RMSD values for all modelled sequences are at around 1.0 Å (with the average of 0.72 and 0.03 standard deviation). The same is true for the A and Z conformations of DNA (see **Figs. S1b,c**).

### Semi-empirical quantum mechanics optimisation of DNA

We further optimised the DNA structures through quantum mechanics (QM) (**Fig. 2b**) by using the PM6-DH+ semi-empirical Hamiltonian under the restricted Hartree-Fock (RHF) approach, as implemented in MOPAC 2016 program^34^. PM6-DH+ with its correction for dispersion interactions, while benefiting from the relative low cost of the semi-empirical QM methods, has successfully reproduced electronic properties of many systems as accurately as the costly QM methods^35^. The water environment was accounted through the intrinsic solvation with Conductor-like Screening Model (COSMO)^36^. The COSMO default 78.4 dielectric constant was used for water, as implemented in MOPAC 2016. For the termination of QM optimisation, we used energy gradient criterion in MOPAC, rather than limiting the optimisation steps. **Fig. S2a1** shows the RMSD of 5’-AAAAAAA-3’ B-DNA from its initial state, as a function of energy gradient cutoff used to optimise the system and look at the structural snapshot. Similar to the MM case discussed above, we found that RMSD plateaus at around 1.0 Å even if a strict convergence criterion is applied. **Fig. S2a2** shows B-DNA structure before and after optimisation until the energy gradient drops below 1.0 kcal/(mol · Å). No substantial conformational change, such as separation of strands, was observed upon such optimisation, keeping the structures within the designated B conformation. On the other hand, 10.0 kcal/(mol · Å) is recommended as the energy gradient criteria for large systems (http://openmopac.net/manual/gnorm.html), such as our heptameric double-stranded DNAs. Therefore, we adopted the maximum gradient of 10.0 kcal/(mol · Å) as our convergence criterion for the QM geometry optimisation. Similar to the MM case, the compliance to low RMSD was observed for all our DNA sequences in their B (**Fig. S2a3**, with the average of 1.00 Å and 0.13 standard deviation). The same was true for A and Z (**Figs. S2b,c**) conformations as well.

### Feature calculation and extraction

We next extracted the electronic and structural features from the obtained refined DNA structures (**Fig. 2c1**). For the electronic features, we conducted further single-point QM calculations on the optimised duplex B, A, and Z-DNA structures. We used the same PM6-DH+ (RHF) with COSMO solvation, but with additional keywords to request more detailed outputs and a full listing of electronic parameters. From these calculations, as general features for each DNA model, heat of formation (Ehof), ionization potential (IP), dipole moment, highest occupied molecular orbital (HOMO) energy, and lowest unoccupied molecular orbital (LUMO) energy were extracted. We also extracted Mulliken charges and populations for the constituent atoms. Since treating charges for all atoms is not realistic for many machine learning setups, we calculated the summarised maximum, minimum, and mean values of the charges and population density values for each of the base, sugar, and phosphate moieties from 1^*st*^ to 7^*th*^ nucleotides at both + and - strands. In the same manner, we extracted electrostatic potential fitted (ESP) charges and populations^37^ and calculated the maximum, minimum, and mean values. Geometric parameters were calculated *via* CURVES+ software^38^, upon which inter- and intra-strand parameters were extracted for the base pair arrangements and backbone angles respectively. Further features were obtained by conducting additional single-point calculations for each strand separately, by masking the other strand in the optimised duplex B-, A-, and Z-DNA structures (**Fig. 2c2**). The difference of Ehof between duplex and the two separate single strands was calculated as an approximate proxy for hybridisation energy. We also considered the duplex state with the 4^*th*^ central base removed and replaced by a hydrogen cap. Single-point calculation was conducted after optimising only this hydrogen position, with a stricter 1.0 kcal/(mol·Å) maximum energy gradient for the convergence criterion. Then, we calculated the difference of Ehof between the complete duplex and the base-removed states, as illustrated in **Fig. 2c2**. This difference should be related to how the central base is stabilised, through the stacking and hydrogen bonding interactions, within the context of the whole DNA sequence.

### Overall calculation costs

For our 24,576 7-mer models, the MM optimisations utilised ≈ 650 hours of CPU time (on average 95.2 seconds per model). The QM optimisations took ≈ 11,052 CPU hours (averaging to 1,618.9 seconds per model). For the subsequent single-point calculations, it took on average 74.1 seconds per model in CPU time. Since there were five such calculations per model, it took overall 2,529 CPU hours. This amounted to 593 CPU days of calculations, which we were able to conduct within about 3 months by utilising up to six computing nodes.

## Data Records

### File description

**Table 1** shows the summary of the database obtained *via* the above procedure. Our database is comprised of 7 deposited dataset files for each k-mer range. The units of features are described in parentheses. The file “energy.txt” includes the overall parameters for the double helical DNA in its B, A and Z states, that is Ehof (kcal/mol), dipole moment (debye), HOMO and LUMO energies (eV), and IP (eV). The file “denergy.txt” includes differences of Ehof upon dehybridisation of the DNA, and the central base removal, calculated for B, A and Z conformations (units are the same as in “energy.txt”). The file “Mullik_Charge.txt” includes the maximum, minimum, and mean Mulliken charge values (in e unites, where e = +1.602177×10^−19^ C) at the base, sugar and phosphate moieties for each nucleotide position in B-, A- and Z-DNA. The file “Mullik_Density.txt” similarly includes the maximum, minimum, and mean values of the Mulliken population density (dimensionless). The files “ESP_Charge.txt” and “ESP_Density.txt” contain datasets for the electrostatic potential fitted charges and populations respectively (units are the same as Mulliken charge and population). The file “Curves.txt” includes intra- and inter-strand geometric parameters for our sequences at their B, A and Z states (Å and degree are the units of distance and angle). The described dataset files include all our sequences, one row of features per sequence, with the first column indicating the sequence in the lexicographic order.

**Table 1.**
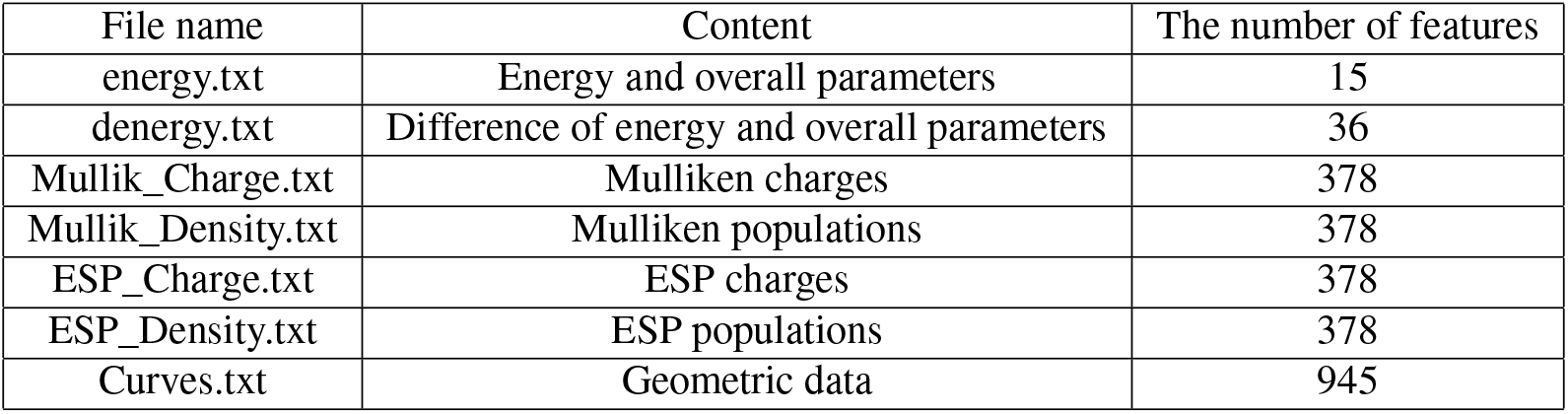
File names, contents, and the number of features of the datasets deposited in the generated database.

| File name | Content | The number of features |
| --- | --- | --- |
| energy.txt | Energy and overall parameters | 15 |
| denergy.txt | Difference of energy and overall parameters | 36 |
| Mullik_Charge.txt | Mulliken charges | 378 |
| Mullik_Density.txt | Mulliken populations | 378 |
| ESP_Charge.txt | ESP charges | 378 |
| ESP_Density.txt | ESP populations | 378 |
| Curves.txt | Geometric data | 945 |

**Table 2** shows our naming rules used to identify the features in the dataset files. Examples are described for (1) energy and difference of energy: B_ds.Ehof means heat of formation energy of duplex B-DNA. B_ds.dEhof_ds_ss means the difference of heat of formation energies between duplex B-DNA and its single-strand states. (2) As an example of charges and populations: here, the plus strand is the strand that has a given sequence, and the minus strand is the complementary strand. For example, when we consider the 5’-AAAAAAA-3’ sequence, the plus strand is the strand that has AAAAAAA nucleotides, while the minus strand is the complementary strand that has TTTTTTT nucleotides. By this rule, B_ds_strandPlus_4_phos_mean.MullikenCharge means the mean value of Mulliken charge of a phosphate part of the 4^*th*^ nucleotide in a plus strand of B-DNA. (3) An example of geometric parameters: B_ds_strandPlus_4.Curves_Xdisp means X displacement of a base of 4^*th*^ nucleotide in a plus strand of B-DNA. Note that the meaning of geometric parameters is well summarised in the 3DNA paper^39^ and **Fig. S3** in this work.

**Table 2.** Explanations of the abbreviations for the feature names used in our database.

| Abbreviation | Explanation |
| --- | --- |
| B_ds, A_ds, Z_ds | Duplex B, A, and Z-DNA. |
| Ehof, Dipole, IP, Ehomo, Elumo | Heat of formation energy, Dipole, Ionization potential, HOMO, LUMO. |
| dEhof_ds_x, dIP_ds_x, dEhomo_ds_x, dElumo_ds_x | Difference of Ehof, IP, Ehomo, and Elumo between the duplex and $x$ states. $x$ = ss(single strand), del1(the centre base at the plus strand is removed), del2(the centre base at the minus strand is removed). |
| strandPlus_x<br>strandMinus_x | $x$-th nucleotide from 5' edge of the strand that has a given sequence.<br>The complementary nucleotide of strandPlus_x. |
| phos_x, sugar_x, base_x | $x$ values for phosphate, sugar, and base parts of a nucleotide. $x$ = mean, max (maximum), min(minimum). |
| MullikCharge<br>MullikDensity<br>ESPCharge<br>ESPDensity | Mulliken charge.<br>Mulliken population.<br>ESP charge.<br>ESP population. |
| Curves_x | Geometric parameters $x$ of DNA calculated by Curves+. $x$ = Shear, Stretch, ... (Intra base parameters), Shift, Slide, ... (inter base parameters), and Alpha, Beta, ... (backbone parameters). To_3 in the inter parameters means inter parameters toward the 3' end side while from_5 means inter parameters from the 5' end side. |

### Characteristics of the DNAkmerQM database

To provide a general exploratory insight into our database, we analysed the loading values^40^ from principal component analysis (PCA)^41^, and describe the characteristics of our datasets. PCA has been utilised to reduce the dimensionality of the datasets to express it through a lesser number of new variables. In such data compression, PCA can output loading values, which are the weights that original variables have for generating new variables. Here, variables that have a similar tendency are likely to have similar loading values. As a result, loading plots, which are obtained by plotting vectors from the origin to loading values of the first principle component (PC1) and the second principle component (PC2), were utilised before for analysing the tendency of variables in biology^41–43^. These plots indicate that variables are positively correlated when their vectors are close to each other and the angle between them is small. As a result, clusters of variables are formed and we may find hidden patterns in the database. By utilizing this, we analyzed relationships between features obtained from our calculations to provide the general insight about the content and interrelation of DNAkmerQM features.

#### Overall loading plots

**Fig. 3** shows loading values from PCA of energy (E), differences of energy (dE), Mulliken charges (MullikC), Mulliken population (MullikD), and geometric features obtained by Curves+ (Curves). Each plot is fitted with an ellipsoid for clarity. **Fig. 3a** shows a loading plot, in which features are grouped by conformations. We found that loadings of B-DNA and A-DNA are along the PC2 direction while Z-DNA is along the PC1 direction. Therefore, B-DNA and A-DNA show a similar tendency while Z-DNA shows a different tendency from them. **Fig. 3b** shows a loading plot grouped by feature types. We found that energy terms (E) remain around the origin of coordinates, while differences in energy terms (dE) lie along the PC2 direction, indicating that they play different roles in overall PCA. On the other hand, loadings for Mulliken charges, Mulliken population, and geometric features overlap with each other, indicating that they have a similar tendency as a whole. **Figs. 3c,d** show a loading plot grouped by the nucleotide locations to which features belong. The features at the 4^*th*^ nucleotide lie along the PC2 direction and they gradually begin to spread along the PC1 direction as they close to the 1^*st*^ nucleotide (**Fig. 3c**). This is also applied to the loading plot for the 4^*th*^ to 7^*th*^ nucleotides (**Fig. 3d**). These indicate that the similarity of features reduces as they close to the edges of DNA.

**Figure 3.**
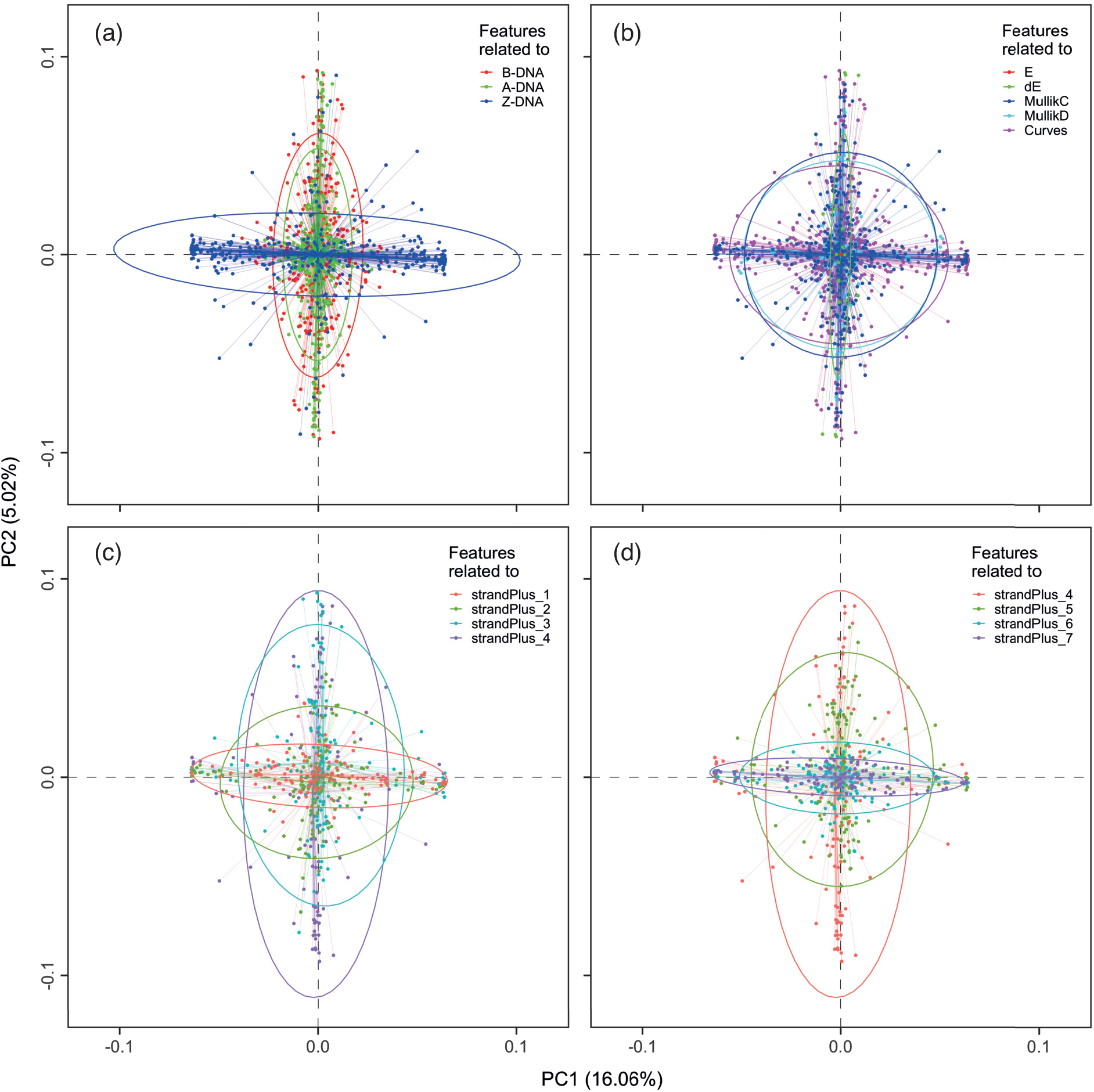
Loading plots from principal component analysis (PCA) of heat of formation energy (E), energy differences (dE), Mulliken charges (MullikC), Mulliken population (MullikD), and geometric (Curves) features. Features are grouped by (a) conformation, (b) feature type, and (c) the positions of nucleotide units at the first half and (d) the second half. The axes represent the first (PC1) and the second (PC2) principal components.

#### Loading plots for energy and energy difference

Here, we conducted PCA for only the features related to energy and differences in energy. **Fig. 4** shows loading values of heat of formation (Ehof) terms obtained by PCA. Colours indicate Ehof of the duplex state, differences in Ehof between the duplex and single-strand states (ds_ss), and differences in Ehof terms between the duplex and the central-base-removed states (ds_del1 and ds_del2). We found that the same kind of Ehof features make clusters, that is, clusters of dEhof_ds_del1, dEhof_ds_del2, and so on. Furthermore, features of B-, A-, and Z-DNA are close to each other in these clusters. These may indicate that DNA conformations have less pronounced effects on these features.

**Figure 4.**
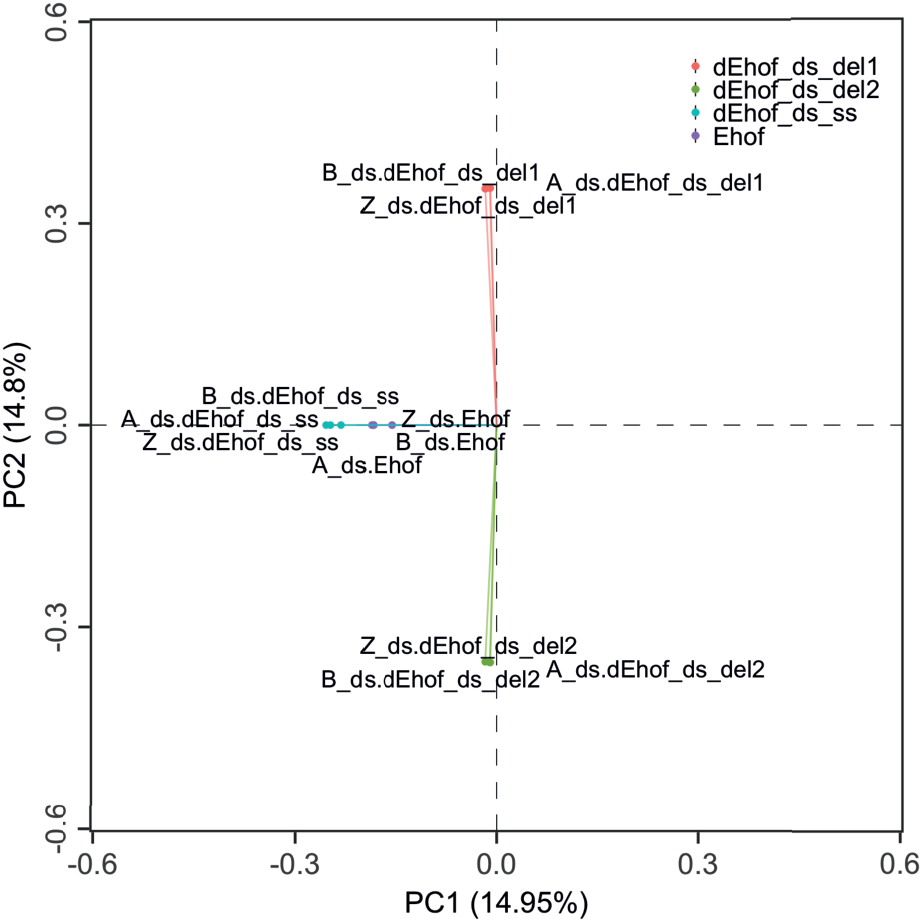
A loading plot for heat of formation (Ehof) terms. Colours indicate energy terms of duplex state (ds), differences in energy terms between the duplex and single-strand states (ds_ss), and differences in energy terms between the duplex and the centre base removed states (ds_del1 and ds_del2).

#### Loading plots for Mulliken charge

**Fig. 5** shows loading plots obtained by PCA from Mulliken charges only. For clarity, we made separate plots for the base, sugar, and phosphate moieties at 3^*rd*^ ∼ 5^*th*^ nucleotide loci. To clearly show tendencies, features are grouped by the value type for base moieties, and by conformation type at sugar and phosphate moieties. We found that, at the base moiety, the maximum values of B-, A- and Z-DNA are clustered together, while the minimum and mean values make a cluster at the StrandPlus_3 position (see the base moieties in **Fig. 5a**). This tendency is also seen at the StrandPlus_4 and 5 positions (base moieties in **Figs. 5b,c**). This indicates that the prevalent characteristic for forming that cluster is the value type rather than conformation. On the other hand, at the sugar moiety, features of B- and A-DNA lie along the PC2 direction, while those of Z-DNA lie along the PC1 direction (sugar moieties of **Figs. 5a-c**). This tendency can also be seen at phosphate moieties (see the phosphate moieties in **Figs. 5a-c**). This indicates that a conformation of DNA is the prevalent characteristic for forming such a cluster at sugar and phosphate moieties, differing from the base.

**Figure 5.**
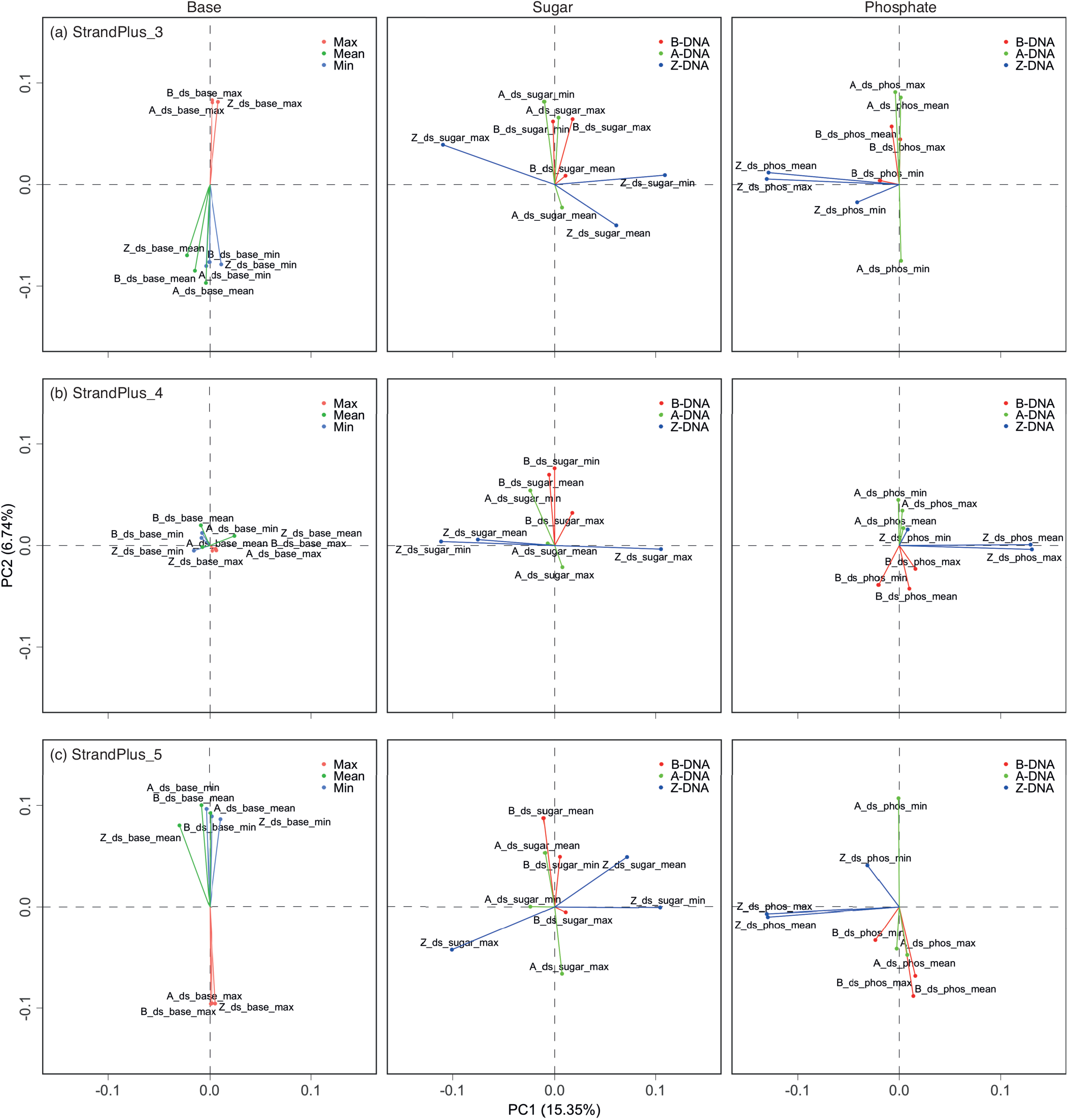
Loading plots from PCA based on only Mulliken charges. Each figure corresponds to the base, sugar, and phosphate moieties at the nucleotide units of (a) strandPlue_3, (b) strandPlue_4, and (c) strandPlue_5 sites. To clearly show tendencies, features are grouped by the value types in the base parts, while features are grouped by conformations in the sugar and phosphate parts.

*Loading plots for geometric parameters*. **Fig. 6** shows loading plots by PCA of only geometric features of B-DNA. For clarity, we made separate plots for the intra, inter, and backbone parts of geometric features at the 3^*rd*^ ∼ 5^*th*^ nucleotide loci. To clearly show tendencies, features are grouped when they show similar behaviours through the 3^*rd*^ ∼ 5^*th*^ nucleotides. We found that the intra parts of geometric features do not make any clusters although Ax-bend, Stagger, and Opening exhibit somehow a similar tendency (see intra parts in **Figs. 6a-c**). On the other hand, H-Twi, H-Ris, Rise, Slide, and Twist make a cluster in the inter parts (**Figs. 6a-c**). Ampli, Phase, Gamma and Zeta for the backbone parts cluster together. Moreover, these features go back and forth between the plus and minus regions of the PC2 axis, similar to H-Twi, H-Ris, Rise, Slide, and Twist in the inter parts. This is also applied to Buckle in the intra parts, with a cluster formed by these features. Similarly, in A-DNA, intra (Shear, Buckle, and Ydisp), inter (H-Twi, H-Ris, Rise, and Roll), and backbone (Alpha, Beta, Zeta, Chi, Gamma, Ampli, and Delta) form a cluster (**Fig. S4**). Z-DNA presents two clusters: a cluster formed by intra (Shear), inter (Roll), and backbone (Alpha, Beta, Chi, and Phase), and a cluster formed only by inter (H-Twi, H-Ris, Twist, Rise, and Slide) and backbone (Epsil, Zeta, Ampli, Delta and Gamma) features, as shown in **Fig. S5**.

**Figure 6.**
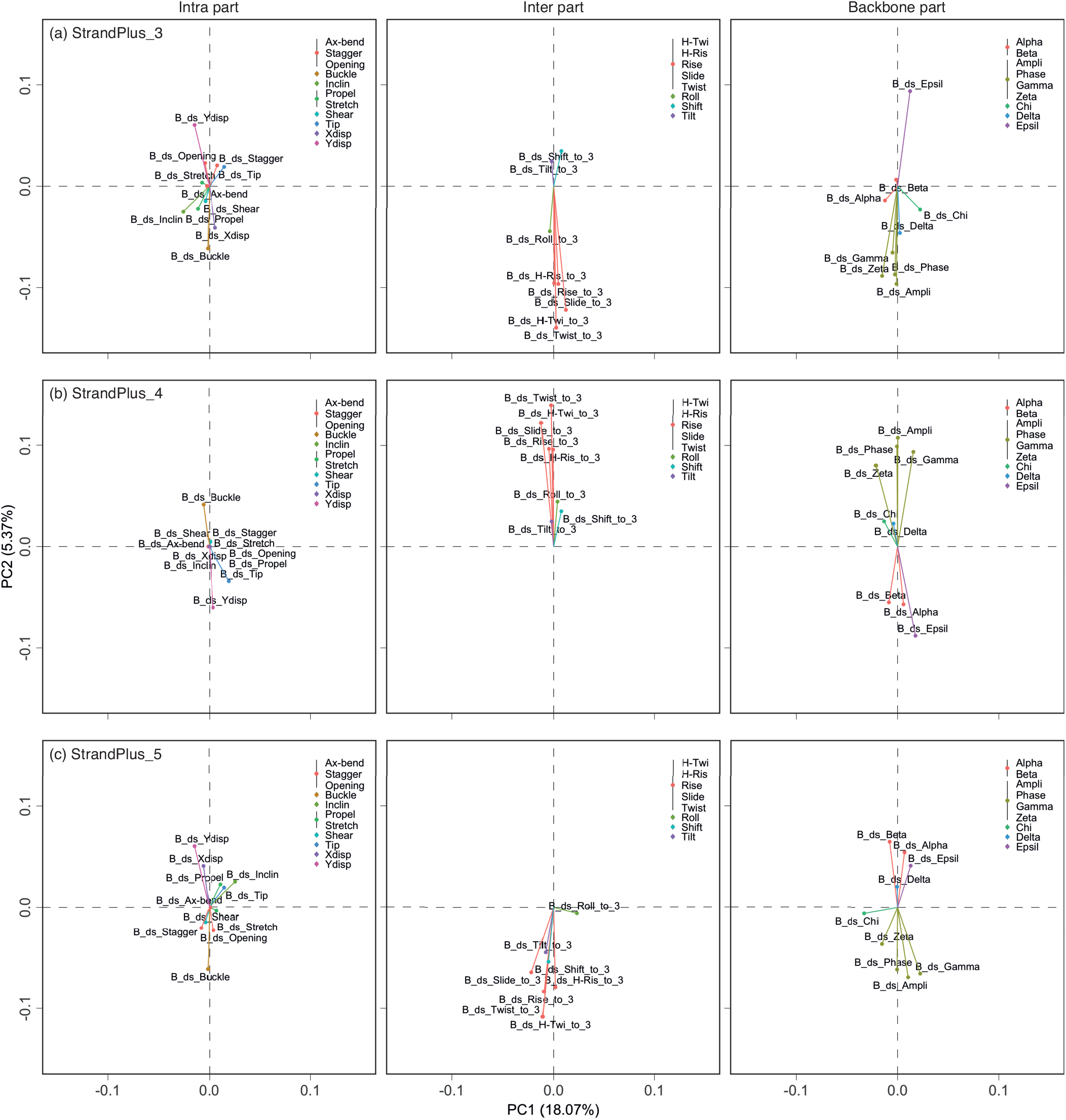
Loading plots from PCA based on only geometric features of B-DNA. Each figure corresponds to the intra, inter, and backbone parts at the nucleotide loci of (a) strandPlue_3, (b) strandPlue_4, and (c) strandPlue_5 sites. To clearly show tendencies, features that show similar behaviours through strandPlue_3~5 are grouped by colours.

#### Correlation analysis for B- and A-DNA

Besides the loading plots, we calculated correlation matrices between features to obtain more insights. **Fig. 7a** shows a heat map for features related to the B-DNA conformation. For clarity, each feature was classified into E&dE, Mulliken Charge, Intra & Inter, and Backbone regions instead of showing all features. Besides the obvious correlations among the features of the same categories (see diagonal parts of **Fig. 7a**), some correlations are also found among features of different categories. For instance, Mulliken charges of base moieties strongly correlate with Zeta and Epsil angles from backbone parameters (**Fig. 7b**), indicating an interesting coupling between the electronic and geometric features in B-DNA. Similarly, we found that Mulliken charges of the phosphate moieties strongly correlate with backbone Alpha in A-DNA (**Fig. S6**).

**Figure 7.**
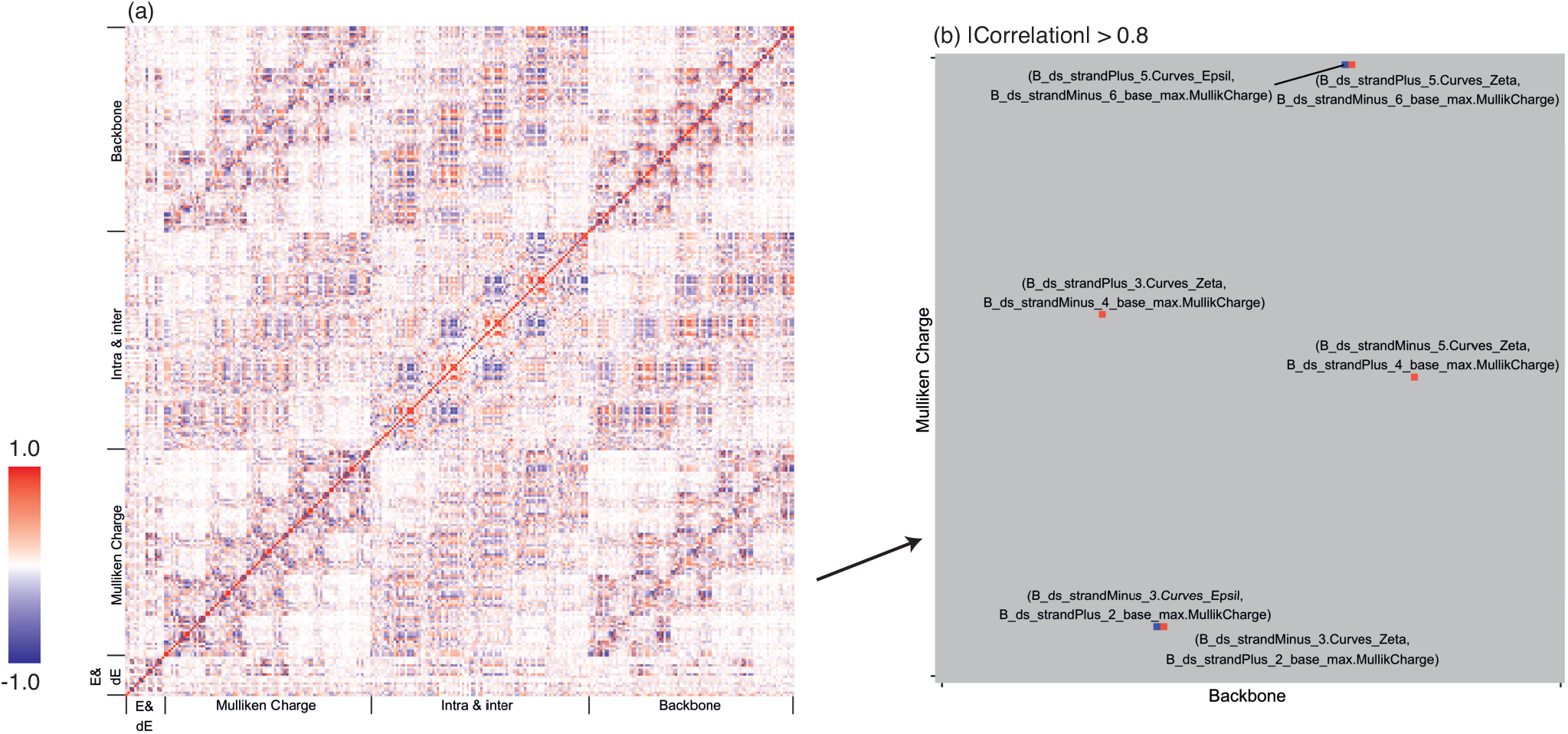
(a) A heat map of features related to B-DNA. For clarity, categories of features are shown instead of showing all features. The colour contour indicates a positive and negative correlation between features. (b) A heat map at the Backbone and Mulliken Charge regions with only high correlation points highlighted.

#### Correlation analysis for Z-DNA

As for the correlation matrix of Z-DNA, a more detailed explanation will be required. **Fig. S7a** shows a heat map for features related to Z-DNA conformation. We found cross correlations all over the plot, which is not observed in B- and A-DNA. Further investigation showed that the Mulliken charge of the phosphate part is strongly correlated with Zeta, Chi, and Delta in backbone parameters. However, strangely, Mulliken charge of phosphate at the 2^*nd*^ nucleotide still correlates with backbone parameters at the 6^*th*^ nucleotide. This demonstrates that parameters away from each other by four nucleotide distance still correlate with each other (a long-range correlation). **Fig. S7c1** shows the mean value of Mulliken charge at the phosphate moieties in Z-DNA. We found that clusters are formed, in which if Mulliken charge is around 0.13, the charge at the next nucleotide is always around 0.16, or vice versa. On the other hand, such a cluster is not observed in B-DNA (**Fig. S7c2**). This indicates that charge values in Z-DNA are not continuous but are seemingly quantised/categorical, i.e. they stay only at around certain values. From these observations, the following can be concluded: The zig-zag backbone structure of Z-DNA interferes with the electronic states in nucleotides, resulting in charge and mechanical values that alternate as small, large, small, large, and so on, values (**Fig. S7c1** right). As a result, long-range order is formed and parameters from nucleotides far away from each other correlate.

## Technical Validation

### Accuracy

Our database and the quality of the features insight reflect the state-of-the art of semi-empirical QM methodology that can still be applied for such a large system and number of molecules. The software and packages we used, that is, AmberTools21, MOPAC, R language and so on have a long history of developments and validations in their respective publications. Furthermore, in the above analyses, we did not find any strange behaviour such as outlier values. Instead, we found that the tendencies of datasets are not so different from our conventional knowledge. For example, the tendencies of datasets for B-DNA and A-DNA (right-handed) are similar but they are different from datasets for Z-DNA (left-handed).

### Applicability

To demonstrate the intended power of our database in actual DNA sequence-based machine learning initiative, below we demonstrate the exclusive use of DNAkmerQM features in predicting context-dependent spontaneous mutation rate constants for A to C mutation *via* machine learning.

### Construction of a dataset for machine learning

The Trek (transposon exposed k-meric mutation rate constants) dataset provides comprehensive sequence-dependent mutation rates for human genome, as obtained from LINE-1 remnants^44^. We combined our QM datasets, which contains features related to B-DNA in “energy.txt”, “denergy.txt”, and “Mullik_Charge.txt” with A to C mutation rate constants (k_A→C_) from the Trek dataset (**Fig. 8a**), resulting in 4096 samples with 102 features.

**Figure 8.**
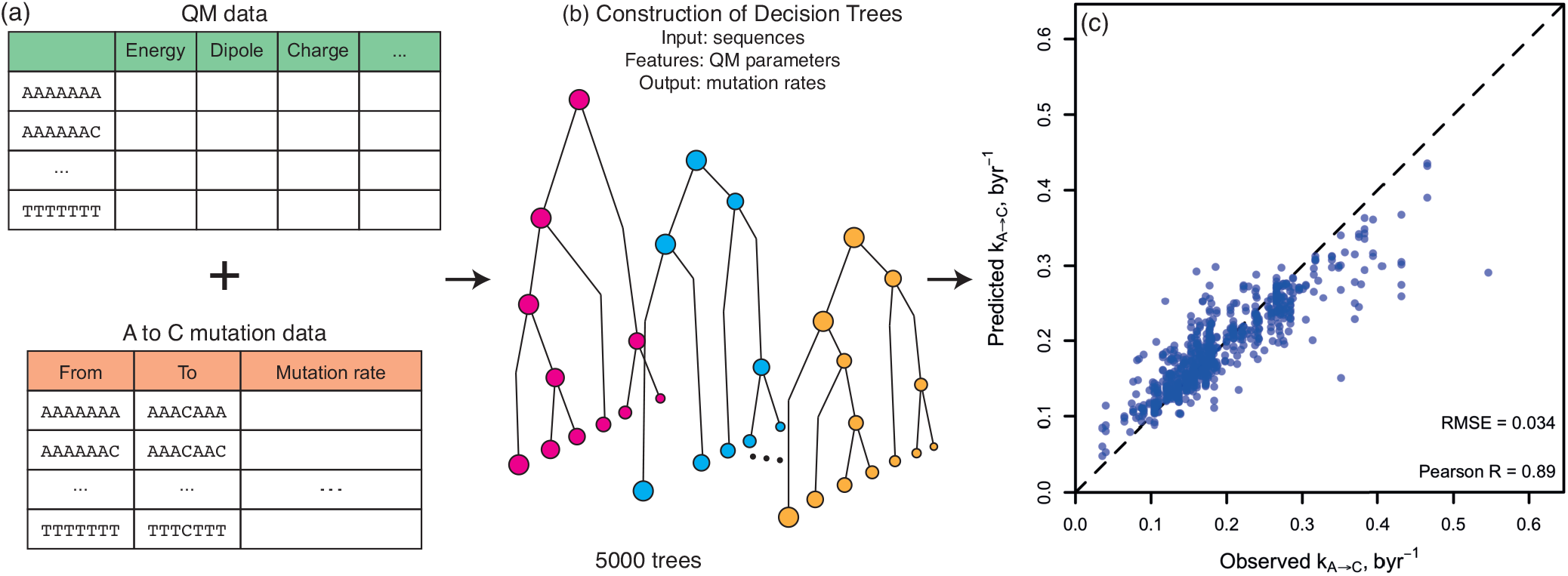
Developing a sequence-driven machine learning model for A to C mutation rates based solely on QM features mapped onto the sequence. Schematic illustrations are shown for (a) combining our QM dataset and an external dataset for A to C mutation rate constants (k_A→C_) and (b) constructing a machine learning model that predicts k_A→C_ from the sequence based on the sequence mapped QM features deposited in the DNAkmerQM database presented in this work. (c) The mutation rate constants predicted through our machine learning model *vs*. actual mutation rate constants (in byr^−1^, i.e. mutation per billion years for each such sequence site). The dashed line shows the diagonal for the ideal match.

### Development of a machine learning model

We divided this dataset into 80% for training (3279 samples) and 20% for pure test (817 samples). Next, we constructed tree-based Gradient Boosting Machine (GBM) models, by which decision trees are consecutively generated to predict the residual values of the ensemble of prior learner trees (**Fig. 8b**)^45–47^. GBMs are known to exhibit superior performance, often prevailing those from neural network-based models for tabulated data^48,49^. GBMs have flexible tunability by five hyperparameters (interaction depth, minimum child weight, bag fraction (sampling rate), learning rate, and the number of trees). These five hyperparameters are related to the overall architecture of the GBM model and drastically affect the performance. The development of GBMs thus iunvolve a careful selection of the optimal combination of its hyperparameters to achieve the best performance. For this, a three-step procedure was employed in this study. (1) Construction of preliminary GBMs by a reasonable initial parameter set. (2) Feature reduction: GBMs provide the importance of features, that is, how much each feature contribute to the performance of GBMs. Based on feature importance values of the preliminary GBM model, we excluded features that do not contribute to the model performance much. This drastically reduced the computation cost of the following procedure. (3) Grid search: after the reduction of features, we developed varying GBMs with various combinations of the five parameters. The performance for each model was measured by the root mean squared error (RMSE) from 10-fold cross-validation. We summarise the employed final hyperparameters (interaction depth = 11, the number of trees = 5000, learning late = 0.01, bag fraction = 0.8, and minimum child weight = 5), along with all the sampled ranges, in **Table S1**.

### Validation of the machine learning model

From the production level GBM model obtained through the above procedure, we predicted A to C mutation rates from the pure test set. **Fig. 8c** shows a dispersion plot between true values in the pure test set and the predicted values by our GBM model. The predicted mutation rate constants agreed very well with the actual values (Pearson’s R = 0.89, RMSE = 0.034). This thus demonstrates the potential of our dataset to provide a wealth of physico-chemical features, which, even while used as sole features, are capable of generating a sophisticated machine learning model for a range of DNA sequence-based biological phenomena.

## Usage Notes

Since the 1^*st*^ and the 7^*th*^ nucleotides are located at the edge of the DNA segments used in the modelling, the usage of their features should be preferentially avoided if used in machine learning. For example, our machine learning model mentioned above shows a lower performance when we include these edge nucleotide data as features. Differences in IP, HOMO, and LUMO (ΔIP, ΔHOMO, and ΔLUMO in “denergy.txt”) do not have clear physical meanings, and thus these features should be avoided too. However, we include them in the database for the estimation of IP, HOMO, and LUMO for single strands and base-deleted states as detailed in the caption of **Fig. 2**.

## Code availability

All software and packages used in this study are freely distributed and available through their citations brought in the text. Our database (https://github.com/SahakyanLab/DNAkmerQM) and the code (https://github.com/SahakyanLab/NucleicAcidsQM) are freely accessible under the LGPLv3 and GPLv3 licenses respectively.

## Acknowledgements

K.M. is supported by JSPS KAKENHI, Grant Number 21J10412 and JSPS Overseas Research Fellowship. A.A.A. is supported by MARA studentship. The Sahakyan Laboratory has been supported by the UK Medical Research Council (MRC), MRC Strategic Alliance Funding (MC-UU-12025).

## Author contributions statement

K.M., A.A.A. and A.B.S. designed the research, K.M. conducted the calculations and analysed the results, A.A.A. contributed analytical techniques to the work, A.B.S. conceived and supervised the research. All authors wrote and reviewed the manuscript.

## Supplementary Information

**Figure S1.**
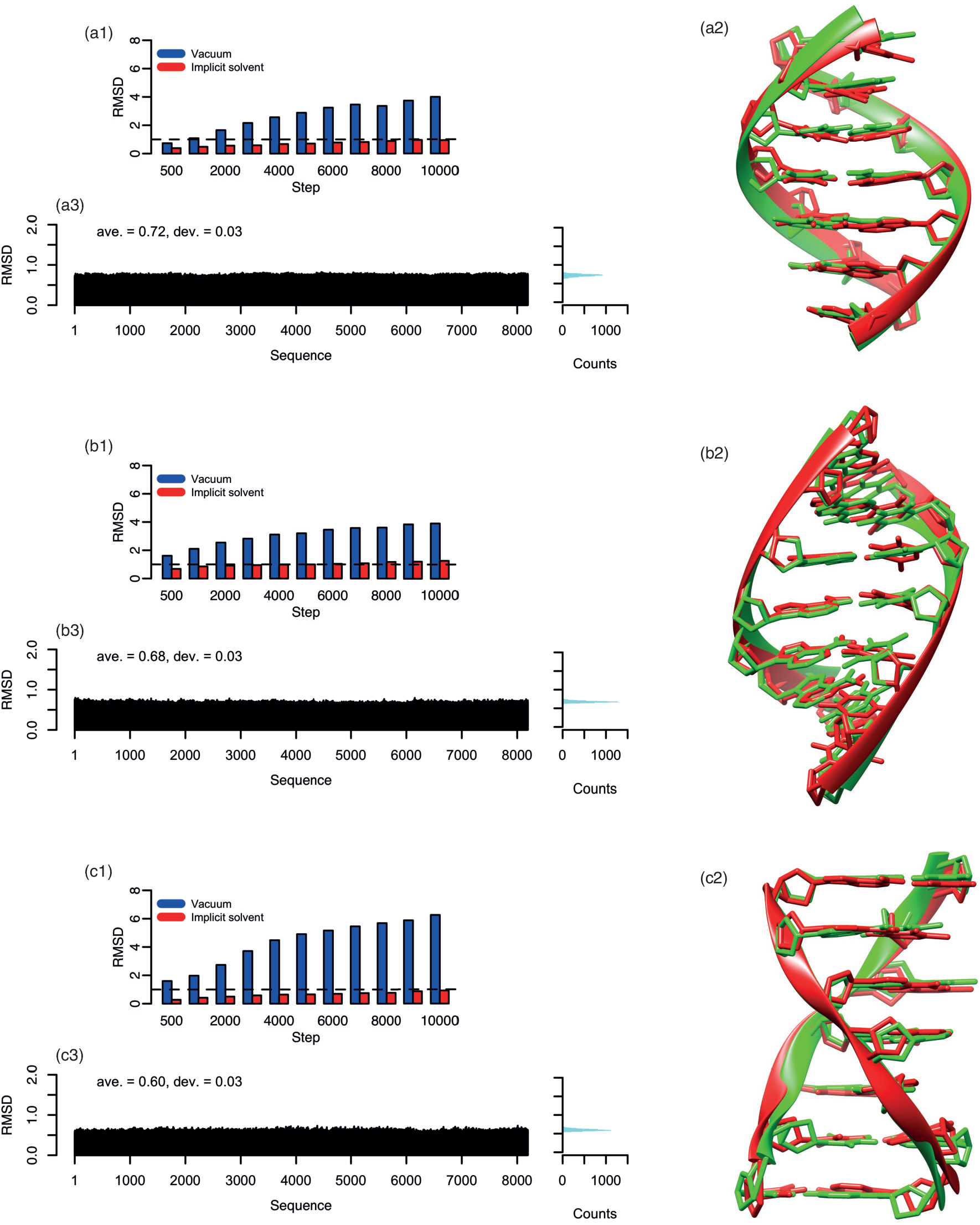
(a1) RMSDs of 5’-AAAAAAA-3’ B-DNA as a function of MM geometry optimisation steps. Red bars indicate RMSDs of the B-DNA in water. For comparison, RMSD of the B-DNA in vacuum is shown by blue bars. (a2) 5’-AAAAAAA-3’ B-DNA structures before (green) and after (red) 5000 optimisation steps. (a3) RMSDs of all 7-mers, 8192 sequences, of B-DNA after 5000 steps of geometry optimisation. Note that numbers were applied to each sequence in a dictionary order. That is, 5’-AAAAAAA-3’ = 1, 5’-AAAAAAC-3’ = 2, 5’-AAAAAAG-3’ = 3, … are indicated. The histogram is attached to the right. Same results for (b) A-DNA and (c) Z-DNA. 2

**Figure S2.**
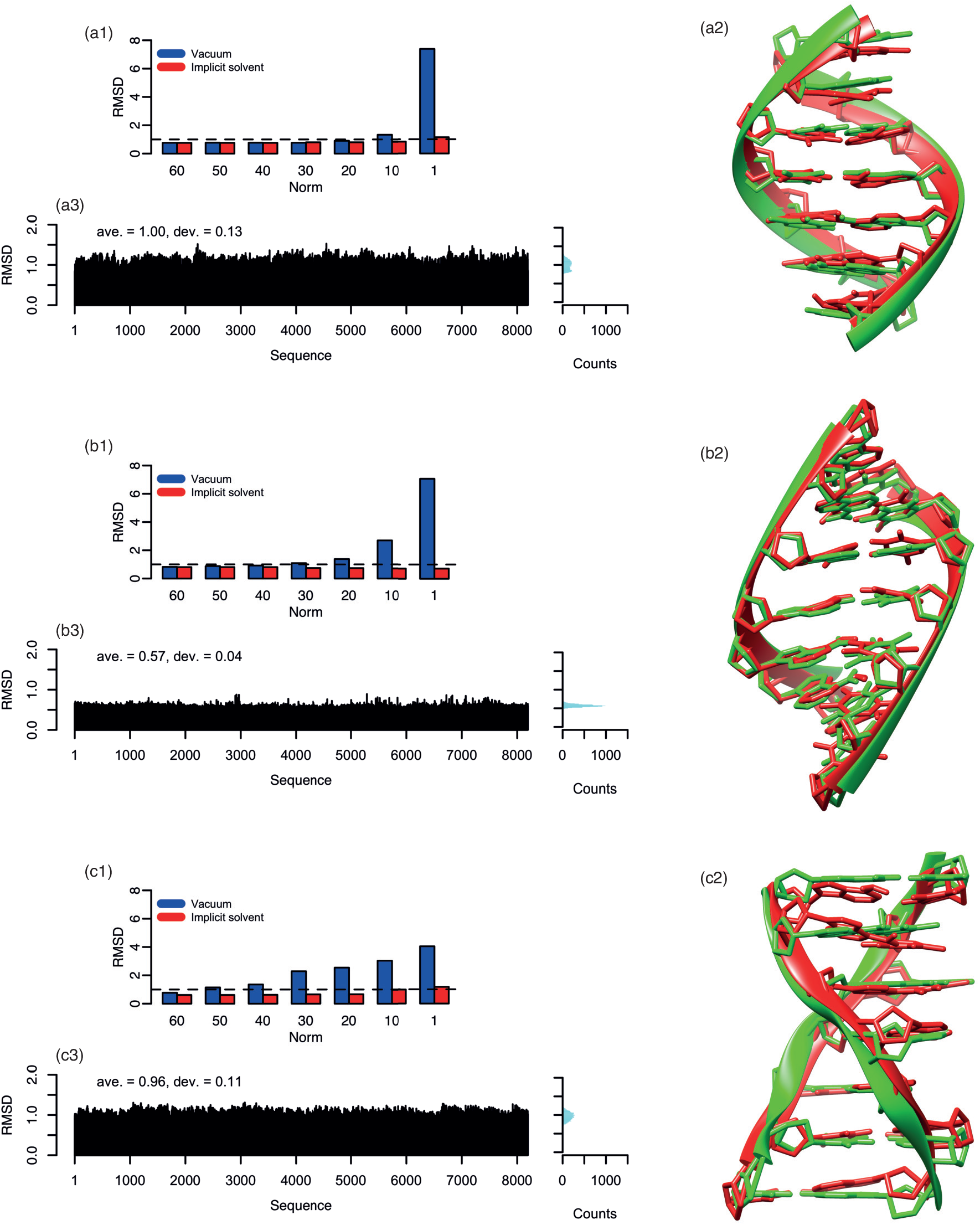
(a1) RMSDs of 5’-AAAAAAA-3’ B-DNA after QM optimisation until an energy gradient norm becomes below a given value. (a2) 5’-AAAAAAA-3’ B-DNA structures before (green) and after (red) optimisation until the corresponding energy gradient norm drops below 1 kcal/(mol · Å). (a3) RMSDs of all 7-mers, 8192 sequences, of B-DNA after optimisation until an energy gradient norm becomes below 1 kcal/(mol · Å). Note that numbers were applied to each sequence in a dictionary order. That is, 5’-AAAAAAA-3’ = 1, 5’-AAAAAAC-3’ = 2, 5’-AAAAAAG-3’ = 3, … are indicated. The histogram is attached to the right. Same results for (b) A-DNA and (c) Z-DNA.

**Figure S3.**
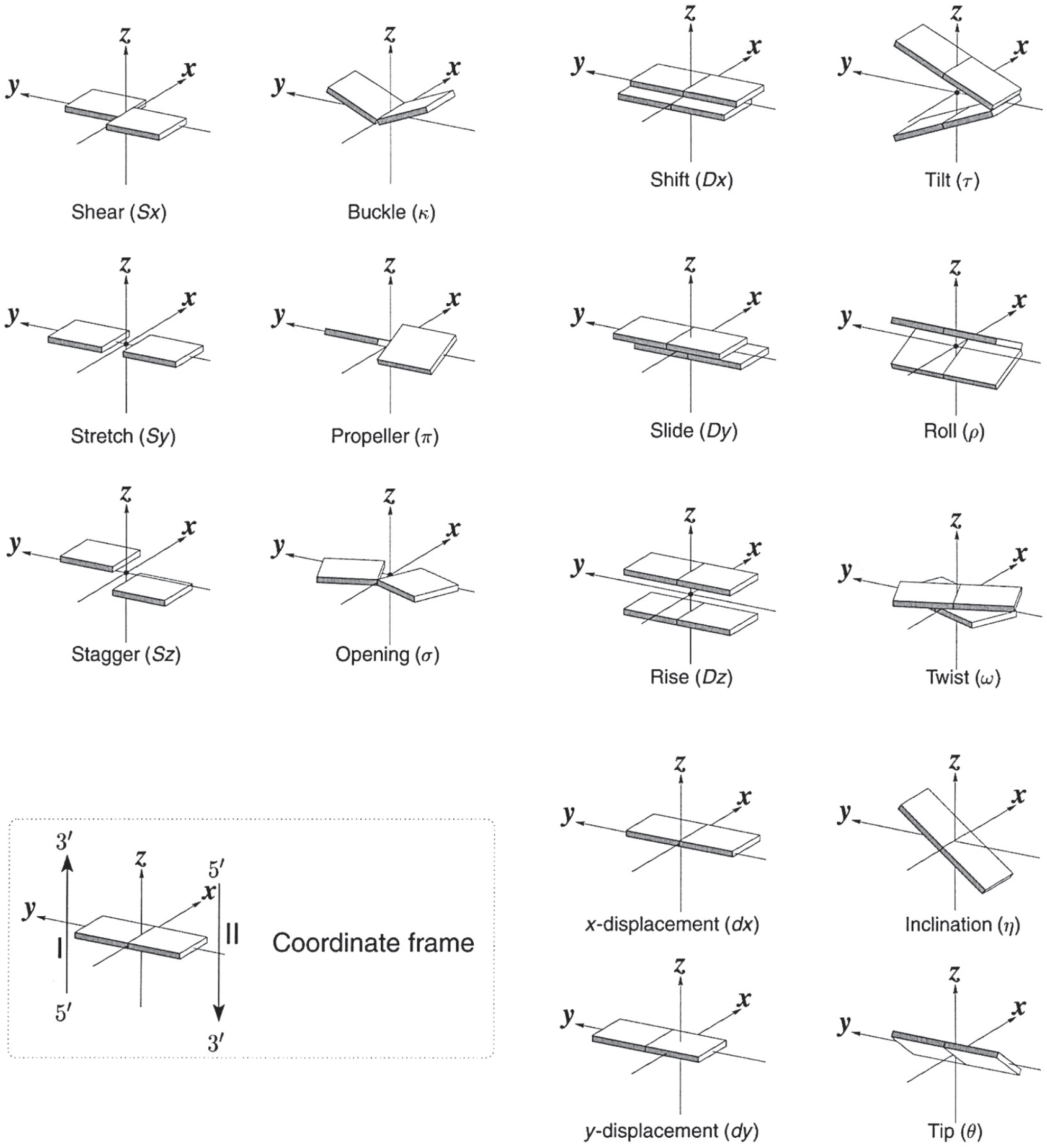
Definitions of the mechanical parameters taken from 3DNA paper (Lu, X. J. and Olson, *Nat .Protoc*. **3**, 1213–1227 (2008))

**Figure S4.**
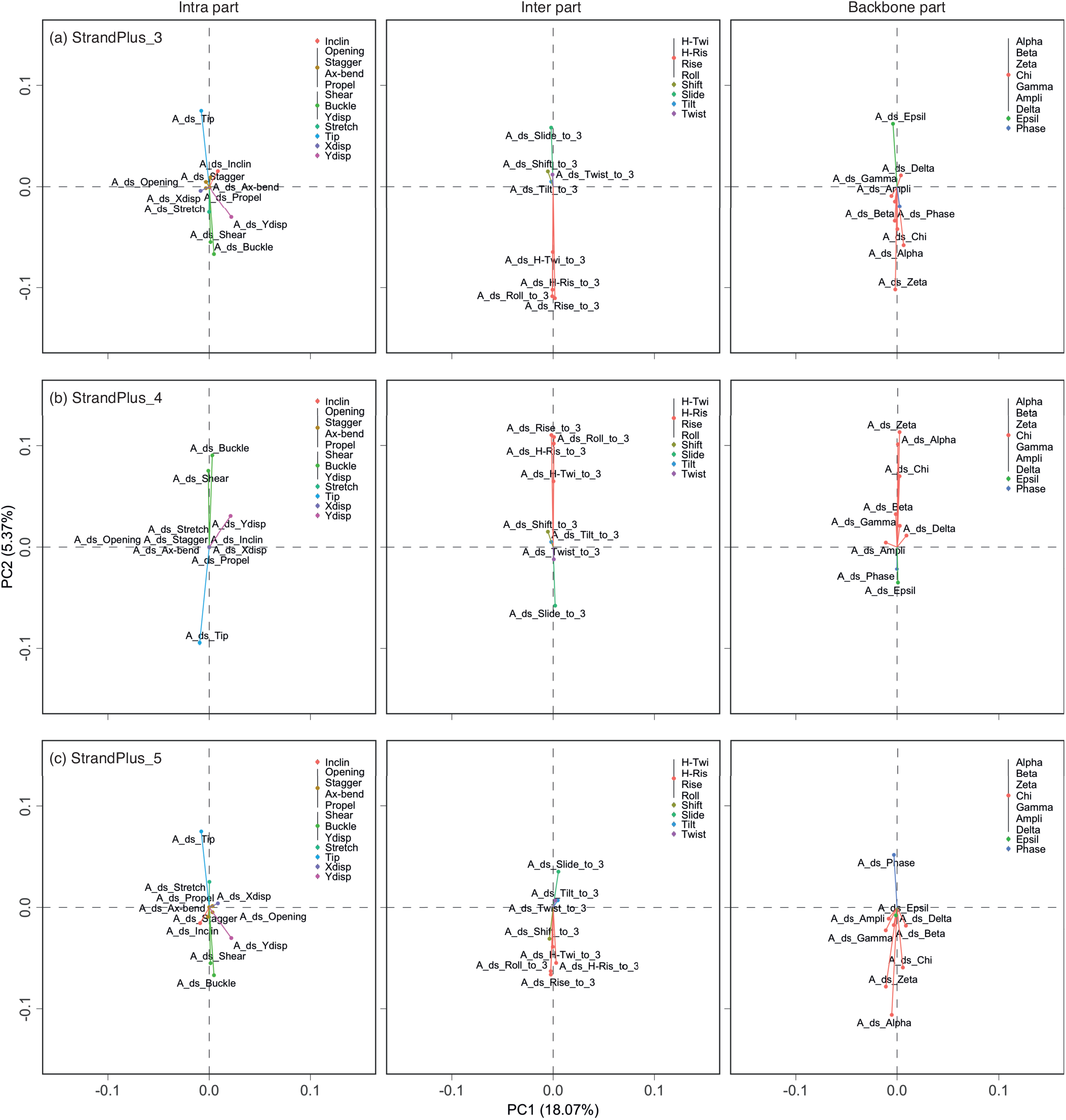
Loading plots by PCA of only mechanical features of A-DNA. Each figure corresponds to the intra, inter, and backbone parts at the nucleotide units of (a) strandPlue 3, (b) strandPlue 4, and (c) strandPlue 5 sites. To clearly show tendencies, features that show similar behaviours through strandPlue 3~5 are grouped by colours.

**Figure S5.**
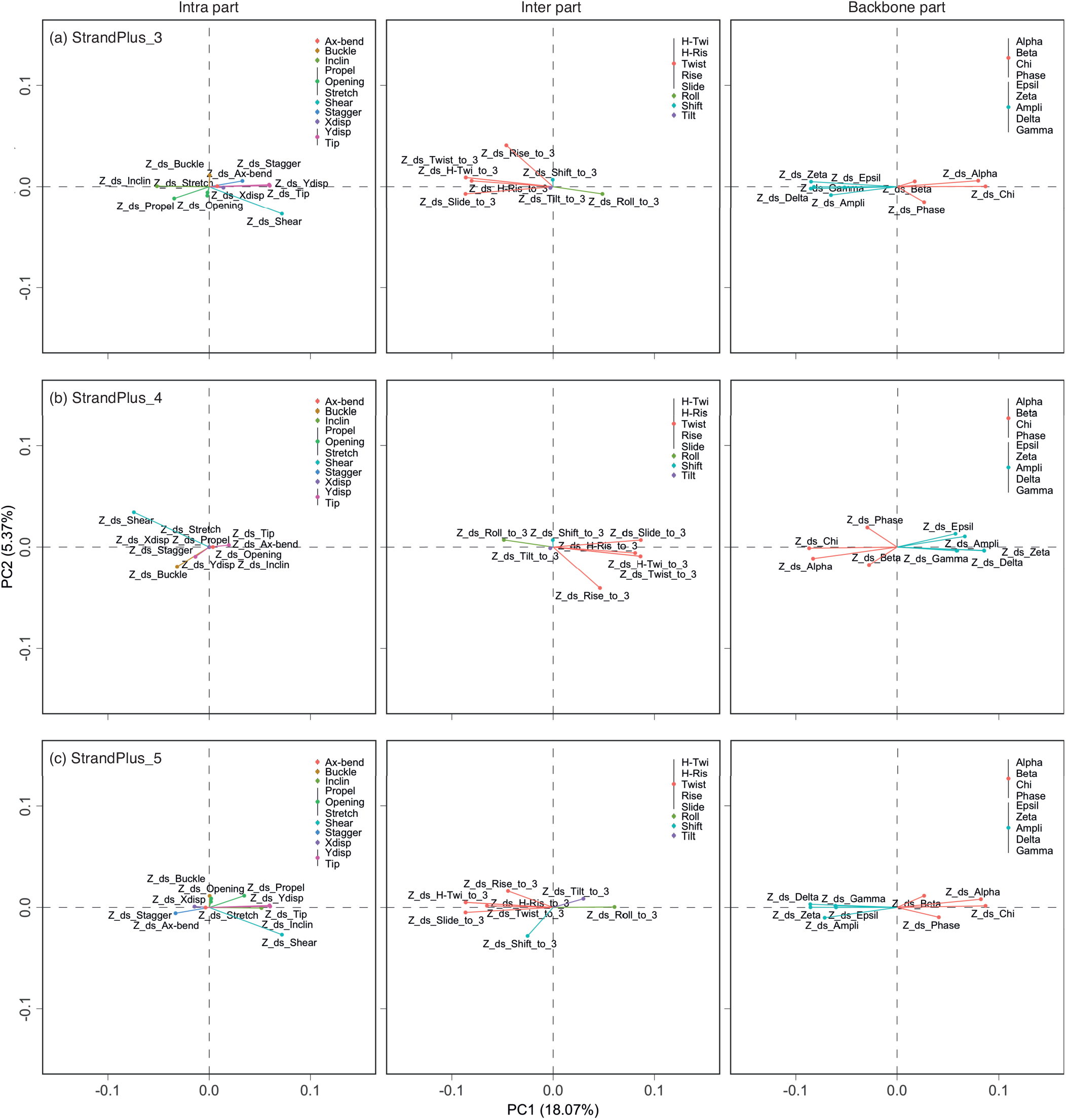
Loading plots by PCA of only mechanical features of Z-DNA. Each figure corresponds to the intra, inter, and backbone parts at the nucleotide units of (a) strandPlue 3, (b) strandPlue 4, and (c) strandPlue 5 sites. To clearly show tendencies, features that show similar behaviours through strandPlue 3~5 are grouped by colours.

**Figure S6.**
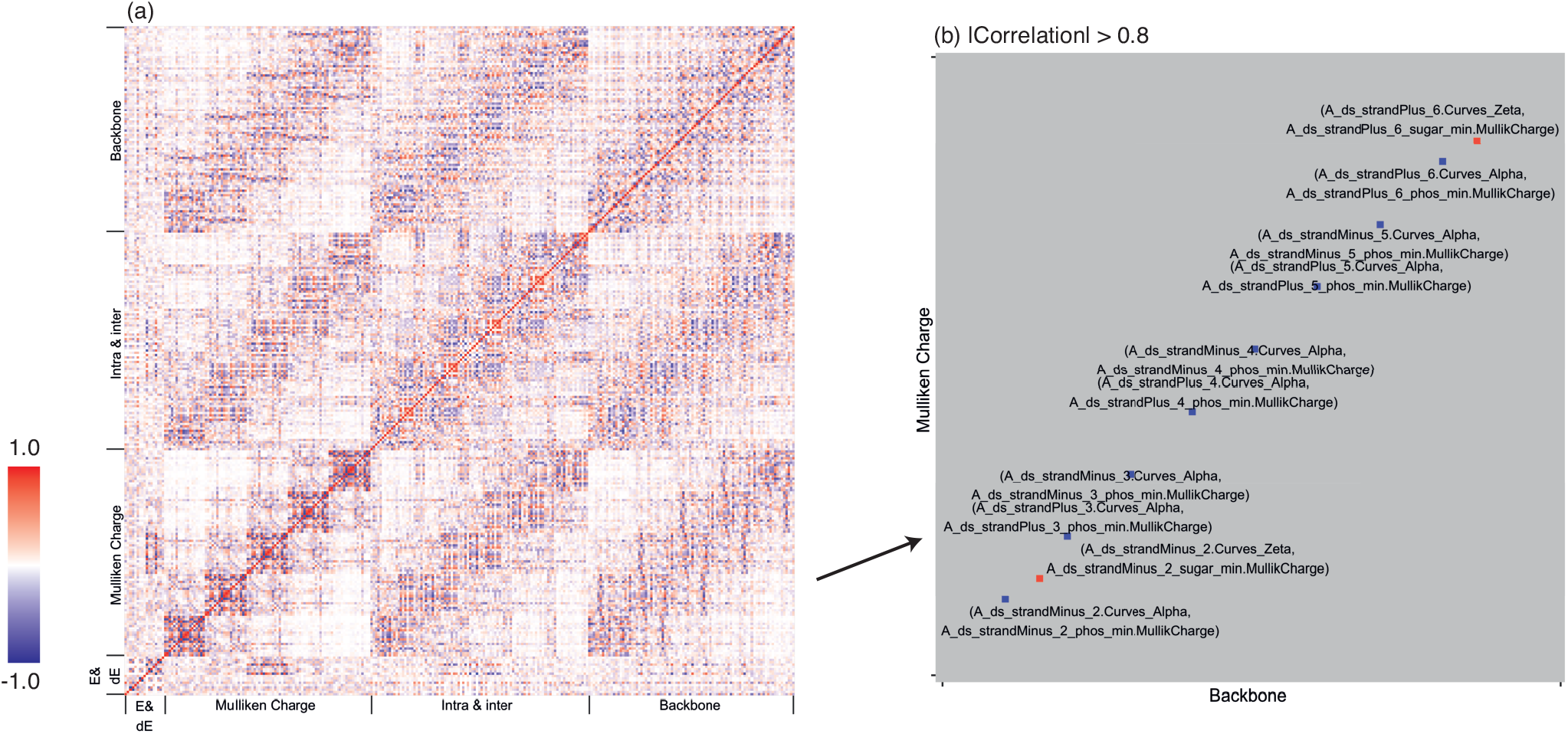
(a) A heat map of features related to A-DNA. For clarity, categories of features are shown instead of showing all features. The colour contour indicates a positive and negative correlation between features. (b) A heat map at the Backbone and Mulliken Charge regions (squares) with only high correlation points being shown.

**Figure S7.**
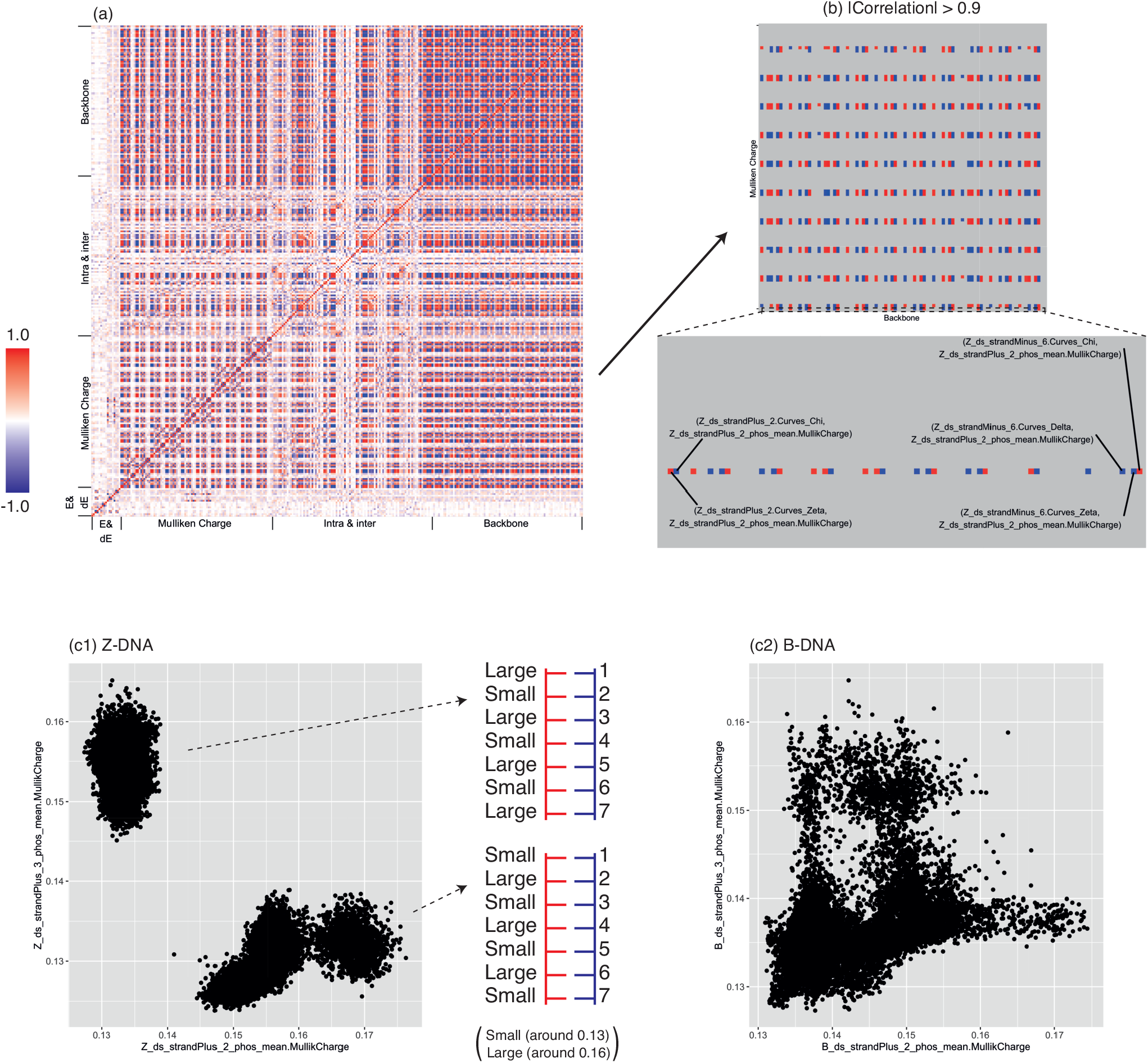
(a) A heat map of features related to Z-DNA. For clarity, categories of features are shown instead of showing all features. Colour contour indicates a positive and negative correlation between features. (b) A heat map at the Backbone and Mulliken Charge square with only high correlation points being shown. A row of Z ds strandPlus 2 phos mean.MullikCharge is enlarged for clarity. (c) The mean values of Mulliken charge of the phosphate parts at the 2^*nd*^ and 3^*rd*^ nucleotides are plotted for Z-DNA and B-DNA. For Z-DNA, schematic images of pseudo-quantised/categorical charge states are shown.

**Table S1.** Hyperparameters for tuning the GBM for A→C mutation rate constants. Values for the architecture that shows the best performance is highlighted in red colour. In feature selection stage, we first generated a model using all features (same as the preliminary model), then modelled using features that have importance > 1 (removing the features that are 100 or more times weaker than the most influential feature in the preliminary model), > 2, > 3, and > 4.

|  |  |
| --- | --- |
| Preliminary GBM model |  |
| Number of features | 102 |
| Number of samples | 3279 |
| Performance metric | RMSE from 10-fold CV |
| Interaction depth | 6 |
| Minimum child weight | 5 |
| Bag fraction | 1 |
| Learning rate | 0.01 |
| Number of trees | {500, 750, 1000, 1500, 2000, 2500, 3000, 3500} |
| Best performance | 0.0423 |
| Feature selection |  |
| Number of features | 102, 75, 47, 35, **29** (Importance>0,1,2,3,4) |
| Number of samples | 3279 |
| Performance metric | RMSE from 10-fold CV |
| Interaction depth | 6 |
| Minimum child weight | 5 |
| Bag fraction | 1 |
| Learning rate | 0.01 |
| Number of trees | {500, 750, 1000, 1500, 2000, 2500, 3000, 3500} |
| Best performance | 0.0387 |
| Rough grid search |  |
| Number of features | 29 |
| Number of samples | 3279 |
| Performance metric | RMSE from 10-fold CV |
| Interaction depth | {6, 9, **12**} |
| Minimum child weight | {1, **5**, 25} |
| Bag fraction | {0.6, **1**} |
| Learning rate | {**0.01**, 0.05} |
| Number of trees | {500, 750, 1000, 1500, 2000, 2500, 3000, **3500**} |
| Best performance | 0.0358 |
| Fine grid search |  |
| Number of features | 29 |
| Number of samples | 3279 |
| Performance metric | RMSE from 10-fold CV |
| Interaction depth | {**11**, 12, 14} |
| Minimum child weight | {**5**, 10 15} |
| Bag fraction | {**0.8**, 1.0} |
| Learning rate | 0.01 |
| Number of trees | {1500, 2000, 2500, 3000, 3500, 4000, 4500, **5000**} |
| Best performance | 0.0354 |

